# Sequence-directed action of RSC remodeler and pioneer factors positions +1 nucleosome to facilitate transcription

**DOI:** 10.1101/266072

**Authors:** Slawomir Kubik, Eoghan O’Duibhir, Wim de Jonge, Stefano Mattarocci, Benjamin Albert, Jean-Luc Falcone, Maria Jessica Bruzzone, Frank C.P. Holstege, David Shore

## Abstract

Accessible chromatin is important for RNA polymerase II recruitment and transcription initiation at eukaryotic promoters. We investigated the mechanistic links between promoter DNA sequence, nucleosome positioning and transcription. Our results indicate that precise positioning of the transcription start site-associated +1 nucleosome in yeast is critical for efficient TBP binding, and is driven by two key factors, the essential chromatin remodeler RSC and a small set of ubiquitous pioneer transcription factors. We find no evidence for recruitment of RSC by pioneer factors, but show instead that the strength and directionality of RSC action on nucleosomes depends upon the arrangement of two specific DNA motifs that promote its binding and nucleosome displacement activity at promoters. Thus, despite their widespread co-localization, RSC and pioneer factors predominantly act independently to generate accessible chromatin. Our results provide insight into how promoter DNA sequence instructs trans-acting factors to control nucleosome architecture and stimulate transcription initiation.

## Highlights

- +1 nucleosome positioning is crucial for TBP binding
- RSC and pioneer transcription factors act independently to position +1 nucleosome
- RSC activity is dictated by the arrangement of two promoter motifs

## Introduction

Protein complexes involved in transcription, replication, repair and recombination need to be in contact with DNA to carry out their functions. Yet in the eukaryotic nucleus the formation of nucleosomes limits access to DNA by its tight wrapping around a histone octamer. A number of factors are believed to be involved in regulating access to DNA: DNA base pair composition, histone post-translational modifications, histone variants, RNA polymerase II (RNAP II), histone chaperones, chromatin remodelers and transcription factors (Lai and Pugh, 2017). However, it is still unclear to what extent each of these factors contributes to nucleosome architecture genomewide, and what rules govern the interplay between them.

Chromatin remodelers are recognized as major players in nucleosome rearrangements that lead to alterations in DNA accessibility. Utilizing the energy of ATP hydrolysis they are able to reposition a nucleosome, change its histone composition, detach DNA from the histone core or even eject the core from DNA (Langst and Manelyte, 2015). Transcription factors have been implicated in recruiting chromatin remodelers to gene promoters (Hartley and Madhani, 2009; Langst and Manelyte, 2015), and histone modifications are known to modulate their activity in vitro (Agalioti et al., 2000; Chatterjee et al., 2011; Ferreira et al., 2007), but detailed mechanistic insight is lacking. Thus the rules governing recruitment and activation of chromatin remodelers at particular genomic nucleosomes still remain elusive.

RSC (Remodels Structure of Chromatin), a member of the highly conserved SWI/SNF subgroup, is the only essential chromatin remodeler in the budding yeast *Saccharomyces cerevisiae.* RSC is able to slide nucleosomes along DNA, either destabilizing or ejecting them (Clapier et al., 2017; Lorch and Kornberg, 2015). Although specific chromatin binding by RSC is difficult to measure by ChIP, there is now general agreement that this remodeler is associated with many gene promoters (Ramachandran et al., 2015). Canonical promoter nucleosome organization is characterized by the presence of a well-positioned (+1) nucleosome that overlaps the transcription start site (TSS) and an upstream (−1) nucleosome (Lieleg et al., 2015). Together, these two nucleosomes demarcate an intervening region which, depending upon its size, will either be nucleosome-depleted (NDR) or accommodate an unstable “fragile nucleosome” (FN) (Kent et al., 2011; Kubik et al., 2015; Weiner et al., 2010; Xi et al., 2011). Functional studies show that RSC contributes to the establishment of promoter chromatin architecture by participating in the generation of NDRs and FNs, since in its absence promoter nucleosome occupancy increases (Badis et al., 2008; Floer et al., 2010; Ganguli et al., 2014; Hartley and Madhani, 2009; Kubik et al., 2015). RSC is specifically implicated in shifting +1 nucleosomes in the direction of the ORF, increasing the size of accessible promoter DNA (Ganguli et al., 2014; Kubik et al., 2015; Yen et al., 2012). Recent genome-wide studies have shown that other chromatin remodelers, in addition to RSC, participate in positioning of the +1 nucleosome (Krietenstein et al., 2016; van Bakel et al., 2013; Yen et al., 2012). Precise positioning of the +1 nucleosome is believed to be important for efficient gene expression (Lomvardas and Thanos, 2001; Rhee and Pugh, 2012) but exactly how this is achieved and how it is mechanistically linked to RNAP II activity are not clear.

RSC affects nucleosome positions at hundreds of promoters genome-wide, but mechanisms underlying its targeting to these promoters are only beginning to emerge. Two out of seventeen subunits of RSC, Rsc3 and Rsc30, bind in vitro to a specific G/C-rich motif (Badis et al., 2008). This motif has been implicated as a site of RSC action in vivo, but a strict relationship between its presence at promoters and the effect of RSC mutation or depletion has not been established (Badis et al., 2008; Ganguli et al., 2014; Kubik et al., 2015). Another DNA sequence feature commonly present at yeast promoters, a polyA tract, was shown to enhance RSC activity in vitro and to promote its action in a specific direction (Krietenstein et al., 2016; Lorch et al., 2014). The role of polyA tracts is also unclear, as they have been proposed either to disfavor nucleosome formation directly (Segal and Widom, 2009; Struhl, 1985) or to stimulate RSC activity (Krietenstein et al., 2016; Lorch et al., 2014). Finally, the functional relationship, if any, between polyA and G/C-rich motifs is unknown. A specific class of TFs, dubbed “general regulatory factors” (GRFs), has also been implicated in RSC recruitment (Hartley and Madhani, 2009). The GRFs resemble vertebrate pioneer transcription factors (PTFs) in being able to “open” chromatin and contribute to the recruitment of co-activators (Zaret and Mango, 2016), and we will thus refer to these ubiquitous yeast TFs as PTFs from hereon. PTFs generally increase DNA accessibility near their binding sites and can act as nucleosome barriers (Badis et al., 2008; Ganapathi et al., 2011; Hartley and Madhani, 2009; Kaplan et al., 2009; Kubik et al., 2015; Mavrich et al., 2008; Yarragudi et al., 2004). At present it is unknown if RSC (i) binds to DNA first and helps to recruit a PTF, (ii) binds to DNA that is made accessible due to PTF binding, (iii) is recruited via direct protein-protein interactions with a PTF, or (iv) is recruited independently of any PTF (Lorch and Kornberg, 2015).

By comparing the changes in genomic nucleosome patterns upon depletion of RSC and three ubiquitous PTFs in budding yeast, we show that both RSC and PTFs exert a global effect on the position of +1 nucleosomes, the majority of which lie just downstream of TATA or TATA-like elements in wild-type cells. RSC, often in conjunction with a PTF, reduces occlusion of these key promoter elements by the +1 nucleosome, and thus favors TBP binding and PIC formation. We present evidence that the strength of RSC remodeling activity is inversely proportional to the distance between paired polyA and G/C-rich motifs, with polyA position and orientation influencing the direction and magnitude of nucleosome movement. Furthermore, we show that the motifs driving RSC action are positioned in a stereotypical manner with respect to PTF binding sites, consistent with functional assays indicating that RSC and PTFs establish nucleosome architecture near the TSS in an independent but coordinated fashion. Our findings thus help to clarify a global mechanism linking specific DNA sequence features to promoter nucleosome architecture and transcription initiation.

## Results

### RSC positioning of the +1 nucleosome strongly influences TBP binding

Although precise positioning of the +1 promoter nucleosome is thought to be important for gene regulation (Nocetti and Whitehouse, 2016; Rhee and Pugh, 2012), mechanistic insight into how this is achieved and its quantitative impact on transcription are still lacking. To address this issue we first investigated the role of RSC, a ubiquitous chromatin remodeler implicated in promoter nucleosome remodeling, paying particular attention to +1 nucleosome position relative to TATA and TATA-like elements. RSC is essential for viability, so to study its activity in vivo we employed the anchor away technique to conditionally and rapidly deplete its catalytic subunit (Sth1) from the nucleus (Haruki et al., 2008; Kubik et al., 2015). We then performed MNase-seq analysis on RSC-depleted and control cells and compared nucleosome positions in each dataset (Kubik et al., 2015). Consistent with previous results (Badis et al., 2008; Ganguli et al., 2014; Hartley and Madhani, 2009; Kubik et al., 2015), we found that RSC depletion caused a general increase in promoter nucleosome occupancy, due largely to upstream shifts of genic nucleosomes, including the +1 nucleosome, and a concomitant downstream movement of the −1 nucleosome (Figure 1A). These shifts were associated with a decrease in nucleosome positioning over gene bodies, as indicated by a pervasive decrease in nucleosome peaks and troughs in these regions. The observed changes were not a consequence of reduced transcription, since depletion of RNAP II has a much weaker effect on nucleosome patterns (Figure S1A).

**Figure 1.**
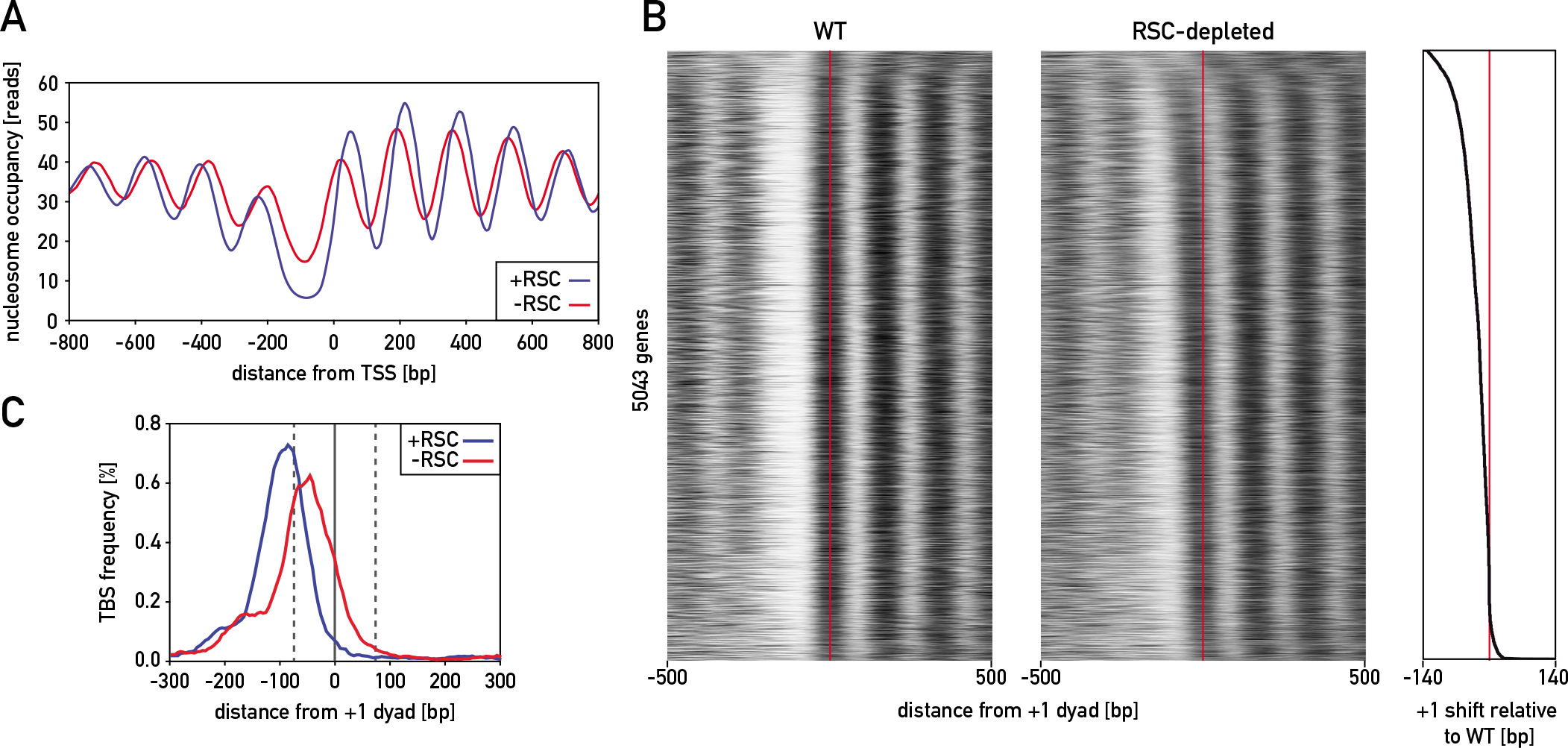
RSC pushes the +1 nucleosome away from the TBP binding site. (A) Average nucleosome occupancy plots, obtained with high MNase concentration, centered on TSS shown in wild-type cells (blue) and cells following nuclear depletion of the RSC catalytic subunit Sth1 (red). (B) Map showing nucleosome positions for 5043 RNAP II-transcribed genes in wild-type cells (left) and Sth1-depleted cells (RSC-depleted; center). Maps were centered on the +1 nucleosome dyad axis measured in wild-type cells (red line) and sorted by the extent of +1 shift (right). (C) Frequency of TBP binding sites (TBS) at genes whose +1 nucleosome significantly changed position after depletion of Sth1 aligned to +1 nucleosome dyad in the presence (blue) and absence (red) of RSC.

To display the effect of RSC at the level of individual promoters, Figure 1B shows nucleosome occupancy maps of 5043 RNAP II-transcribed genes (rows), centered on the +1 nucleosome dyad axis and sorted by the magnitude of the +1 shift caused by RSC removal, for control and RSC-depleted strains. The diagram to the right of the two maps (Figure 1B) plots the magnitude and direction of the +1 nucleosome shifts and shows that for nearly 70% of all genes movement is in the upstream direction when RSC is depleted, with the +1 nucleosome encroaching the NDR. Notably, at those promoters where the upstream +1 shift was >25 bp their TATA or TATA-like elements (referred to hereon as TBP binding sites, TBSs) are mostly located upstream of or just within the DNA exit point of the +1 nucleosome in the presence of RSC (Rhee and Pugh, 2012) but become predominantly nucleosome-associated upon RSC depletion (Figure 1C). In contrast to TBSs, the motifs proposed to recruit or stimulate RSC (Badis et al., 2008; Krietenstein et al., 2016; Kubik et al., 2015; Lorch et al., 2014) mostly remain outside of the +1 nucleosome boundary, and are thus presumably accessible even in the absence of RSC action (Figure S1B and C).

Encroachment of the +1 nucleosome over the TBS following RSC depletion implies a direct effect on PIC assembly and transcription. To test this possibility we performed ChIP-seq of both Myc-tagged TBP (Spt15) and RNAP II following Sth1 nuclear depletion. We observed a decrease in TBP signal after depletion of Sth1, particularly at promoters whose +1 nucleosome was strongly shifted upstream (examples shown in Figure 2A). Indeed, genome-wide nucleosome occupancy changes measured at nearly 6000 TBSs (Rhee and Pugh, 2012) were strongly correlated with changes in TBP binding at these sites (Pearson R=-0.44, p<10^−6^; Figure 2B). Amongst the TBSs associated with RNAP II-transcribed genes, we found 1364 such sites where the TBP signal decreased at least 1.5- fold (average plots in Figure 2C). This decrease in TBP binding was associated with diminished RNAP II levels at the corresponding gene bodies (average plots in Figure 2D; individual examples in Figures 2A) and the correlation between the two factors was strong (Pearson R=0.65, p<10^−7^) (Figure 2E). The simplest explanation of the above data is that RSC shifts the +1 nucleosome away from the TBS at the majority of RNAP II-transcribed genes, which in turn stimulates PIC formation and transcription. In accord with published results (Tramantano et al., 2016) we observed no major changes in nucleosome patterns at promoters of the vast majority of genes upon TBP depletion (Figure S2A) except for tRNAs (Figure S2B) and a small number of genes which display high levels of TBP (Figure S2C and S2D). We conclude that TBP contributes to +1 nucleosome positioning only at a limited number of targets characterized by remarkably strong TBP binding (de Jonge et al., 2017).

**Figure 2.**
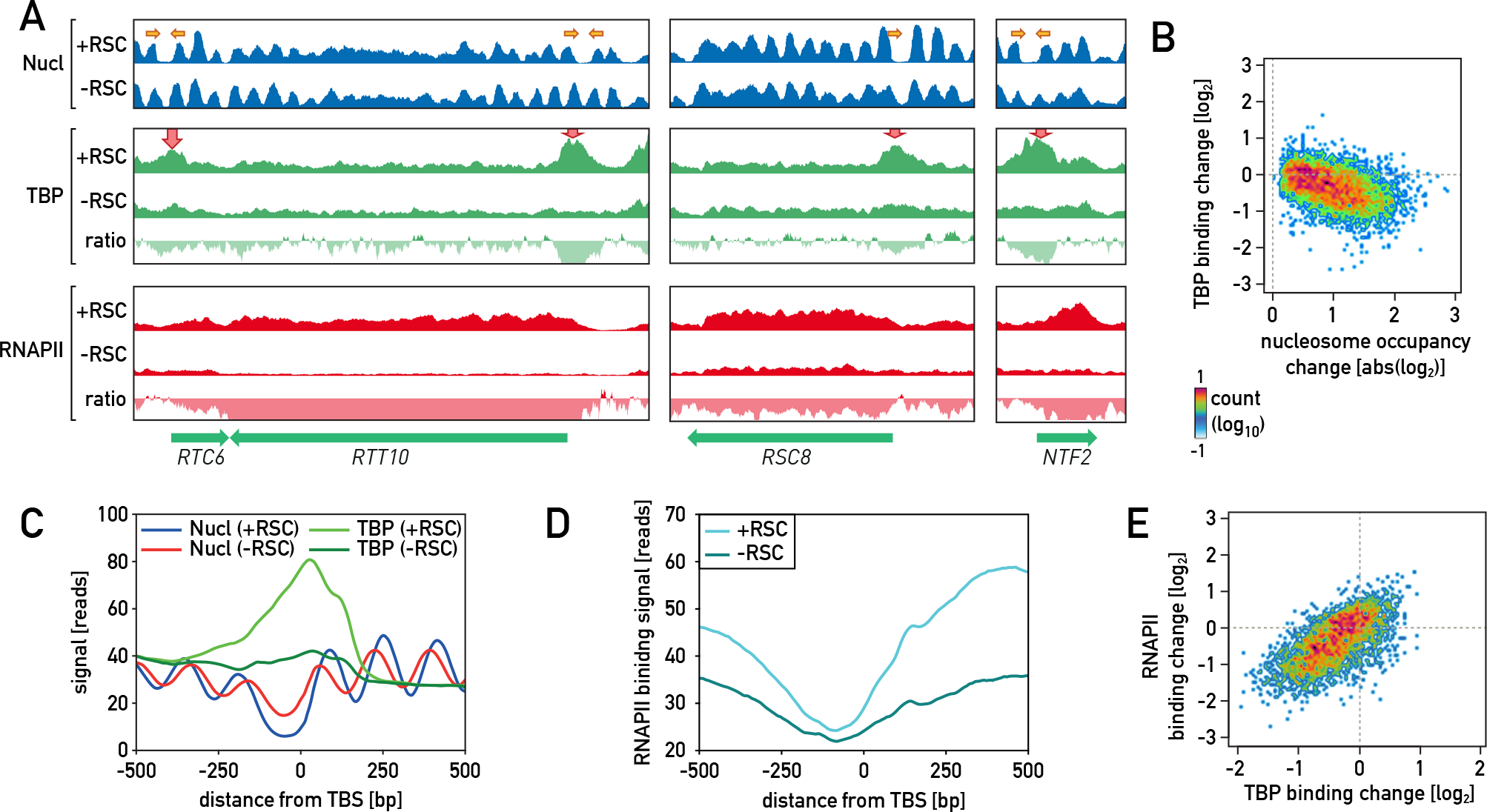
RSC facilitates binding of TBP. (A) Snapshots of the three genomic regions with tracks showing, for wild-type and RSC-depleted cells (+/-RSC): nucleosome occupancy, TBP ChIP-seq signal and signal ratio (-RSC/+RSC), RNAP II ChIP-seq signal and signal ratio (-RSC/+RSC). Arrows point to promoters where nucleosome changes upon RSC depletion are accompanied by decrease in TBP and RNAP II signals. (B) Scatterplot showing the relationship between absolute nucleosome occupancy change upon Sth1 depletion and TBP binding change calculated in 200 bp-wide regions centered on all TBSs (Rhee et al., 2012). (C) Plot showing nucleosome occupancy and TBP signals, in the presence and absence of RSC, centered on protein coding gene-associated TBSs displaying at least 1.5-fold decrease in TBP signal upon RSC depletion. (D) Plot showing RNA Pol II signals in the presence and absence of RSC at the same regions as in (C). (E) Scatterplot showing the relationship between TBP binding change calculated in 200 bp-wide regions centered on TBSs associated with RNA Pol II genes and RNA Pol II binding change in the ORF of these genes.

### DNA sequence motif proximity and polyA tract orientation regulate RSC binding and activity

The data presented above indicate that RSC increases the accessibility of the TATA element for TBP binding, but do not explain how the remodeler’s activity is targeted towards particular promoter nucleosomes or how its activity is regulated. Since previous studies indicated a key role of polyA and G/C-rich motifs in RSC action (Badis et al., 2008; Hartley and Madhani, 2009; Krietenstein et al., 2016; Kubik et al., 2015; Lorch et al., 2014), we decided to examine more carefully the distribution and orientation of these motifs at promoters genome-wide. We began with the assumption that the motifs work together and searched all promoters for motif pairs. To simplify the analysis, we only considered motif pairs where for a given polyA motif the paired G/C-rich motif was the closest such motif, and vice versa. Next, we divided the motif pairs (3205 in total) into 4 groups based on motif orientation, sorted the members in each group by the distance between the motifs, and centered the resulting maps on the midpoint between the paired motifs (Figure 3A, left). All groups were represented approximately equally, arguing against a selection for any specific motif orientation.

**Figure 3.**
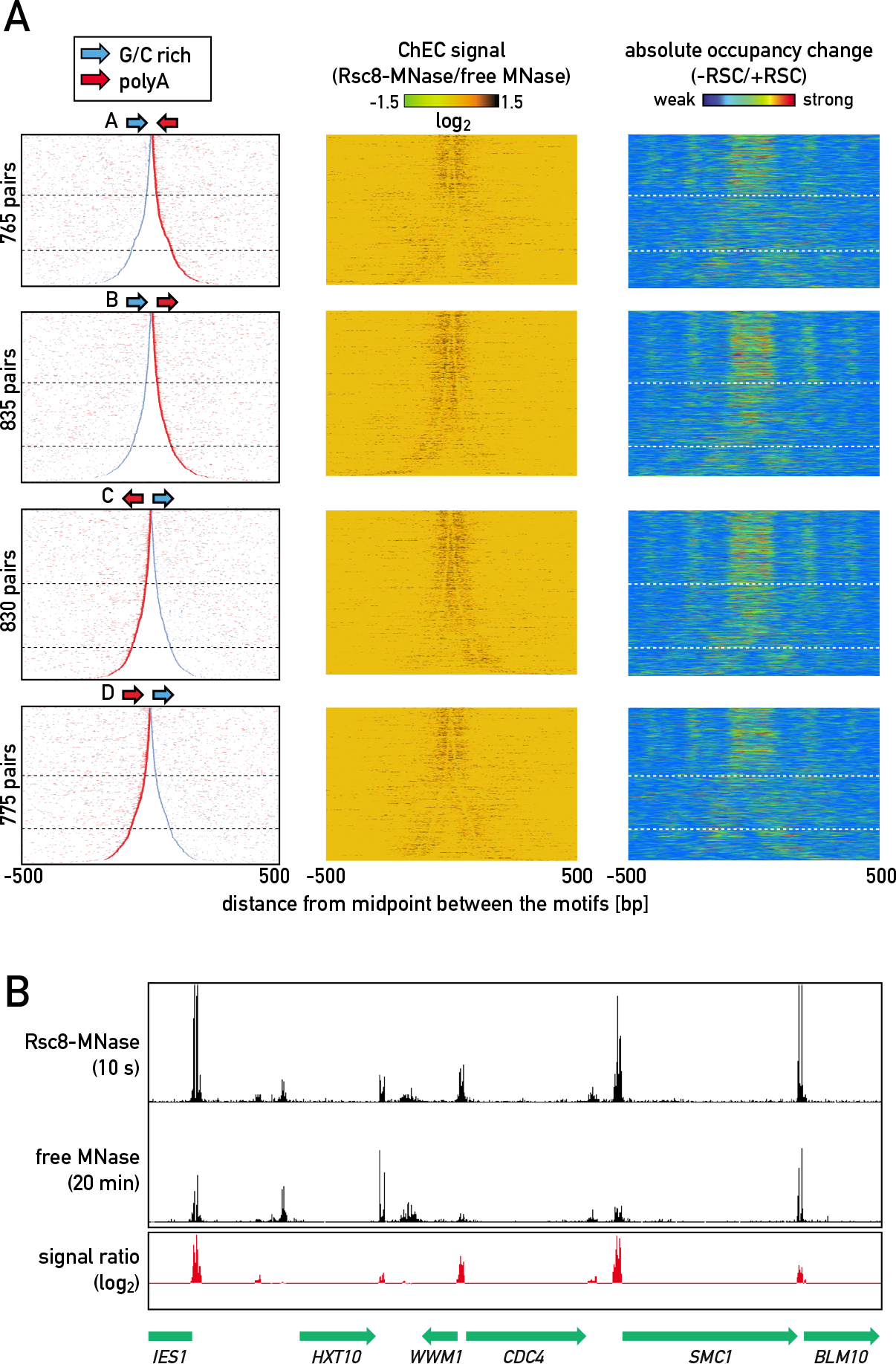
ChEC-seq reveals RSC binding at the polyA and G/C-rich motifs. (A) Maps showing promoter regions with 4 possible motif orientations (A-D), sorted by intermotif distance (left), RSC ChEC signal (normalized to free MNase) in the same regions (middle) and absolute nucleosome occupancy change upon depletion of RSC (right); vertical dashed lines indicate boundaries between motifs positioned close (distance between the motifs <40 bp), separated (40 – 150 bp) and isolated (>150 bp). (B) Representative fragment of the genome with ChEC signal of Rsc8-MNase, free MNase and signal ratio (smoothed by 5 bp).

We next asked how motif spacing and orientation is related to RSC binding. Since ChIP of chromatin remodelers, including RSC, is notoriously inefficient (Yen et al., 2012), we turned to the Chromatin Endogenous Cleavage (ChEC) assay to measure RSC binding (Schmid et al., 2004), using deep sequencing to quantify sites of MNase cleavage by a chromosomally encoded Rsc8-MNase fusion protein (ChEC-seq; (Zentner et al., 2015); see STAR Methods for details). As a control for non-specific MNase cleavage we used a strain expressing “free” MNase from the *REB1* promoter (Zentner et al., 2015). The general profiles (peak locations) of the Rsc8-MNase and free MNase digests were similar, with cleavage sites primarily mapping to intergenic regions, suggesting that both Rsc8-MNase and free MNase can sample “open” chromatin at promoters and terminators (see Figure 3B for a representative example). Nevertheless, many sites were cleaved by Rsc8-MNase more readily than the free MNase control. To obtain a conservative estimate of specific Rsc8 binding we calculated the ratio of read counts in peaks from an early time point of Rsc8-MNase cleavage (10 sec) to those of a free-MNase digest that yielded a maximal signal genome-wide (20 min). In support of the validity of our Rsc8 ChEC analysis, we found that the calculated Rsc8-MNase signal was strongly correlated with the measured nucleosome occupancy change at +1 nucleosomes upon RSC depletion, whereas signal strength in free MNase digests showed no such correlation (Figure S3A).

We next mapped Rsc8-MNase/free MNase read count ratios onto the motif-containing promoter regions defined above. This revealed a striking correlation between RSC binding affinity and motif proximity (ordered from top to bottom; Figure 3A middle). Significantly, a similar relationship was observed between motif proximity and nucleosome occupancy change upon RSC depletion (Sth1 anchoring; Figure 3A right). The correlation between protein binding signal and nucleosome changes upon Sth1 depletion indicates that ChEC-seq of Rsc8 reflects the binding of RSC globally significantly better than either native ChIP-seq of Sth1 (Ramachandran et al., 2015) or standard ChIP-seq of Rsc8 (Yen et al., 2012) (Figure S3B). Consistent with our previous results (Kubik et al., 2015) RSC binding (Rsc8 ChEC-seq) and nucleosome occupancy changes upon depletion of this remodeler were associated globally with the abundance of polyA and G/C-rich motifs in promoter regions (Figure S4A, S4B and S4C). Taken together, these data reveal that the proximity of paired polyA and G/C-rich motifs promotes RSC binding and nucleosome displacement.

To investigate possible effects of motif orientation on RSC binding we aligned the ChEC-seq data for Rsc8-MNase to either the polyA or G/C-rich motif in the four different groups. Significantly, alignment to the 5’ end of the polyA tract revealed a notable peak of ChEC signal precisely at this site, but only in groups B and C where the G/C-rich motif is located 5’ of the polyA tract (Figure 4A). In contrast, we detected little if any orientation bias in Rsc8 ChEC signal relative to the G/C-rich motif (Figure 4B). Instead, though, there was a pronounced absence of signal at the G/C-rich motif itself in all four classes possibly due to protection by bound Rsc3/Rsc30 subunits. A detailed examination of nucleosome occupancy change upon RSC depletion as a function of motif position and orientation also revealed a stronger effect upstream of the polyA motif. This effect was observed for all classes when the motifs were separated or isolated (n>40 bp) but most pronounced at separated motifs of groups B and C, where the G/C motif is located 5’ of the polyA (Figure 4C). In summary, these data show how specific promoter DNA sequences direct the binding and/or activity of RSC.

**Figure 4.**
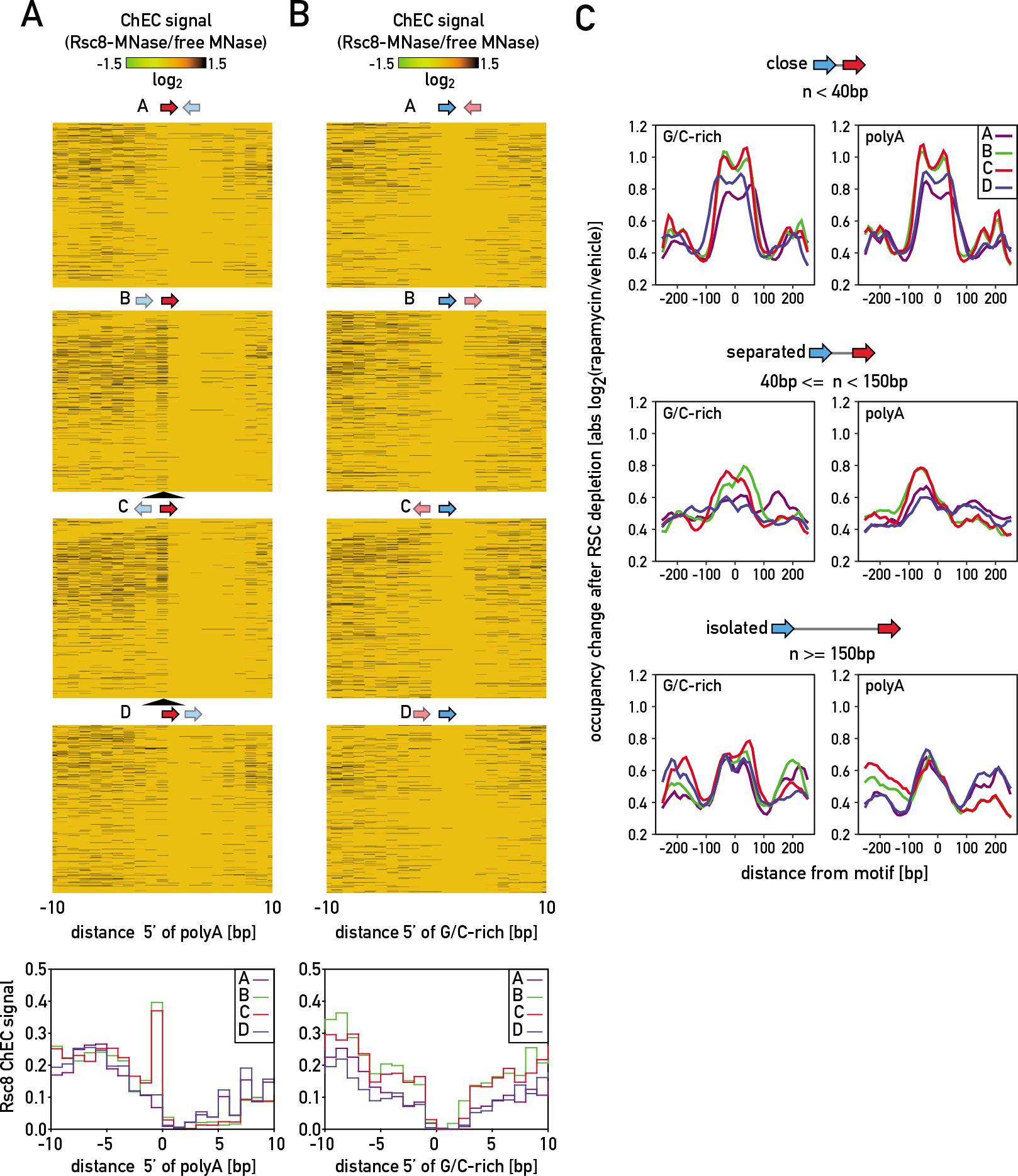
Motif arrangement regulates RSC binding and activity. (A) Heatmap of Rsc8 ChEC signal centered on 5’ side of polyA motifs sorted like in Figure 3A and the average signal plots (bottom). (B) Rsc8 ChEC signal as in (F) but centered on G/C-rich motifs. (C) Average absolute nucleosome occupancy change, centered on G/C-rich (left) or polyA (right) motifs, categorized according to motif-motif orientation (A-D) and distance (top to bottom).

### PTFs and RSC act independently but coordinately at most promoters

Chromatin remodelers such as RSC are not the only protein factors involved in shaping the promoter nucleosome landscape. PTFs bind at many promoters, where they also contribute to the formation of NDRs or FNs immediately upstream of the TSS-associated +1 nucleosome (Ganapathi et al., 2011; Hartley and Madhani, 2009; Kubik et al., 2015). Both PTFs and RSC often act at the same promoters, but, as pointed out above, the functional relationships between the two are unresolved.

To address this issue we used the anchor-away method to simultaneously deplete from the nucleus the RSC catalytic subunit Sth1 and either of three ubiquitous PTFs: Abf1, Reb1 or Rap1. The effects of these double-depletion experiments were then compared to depletion of each factor alone for every promoter PTF motif found within 400 bp of a TSS for which binding was confirmed by ChIP-seq analysis (Kasinathan et al., 2014; Knight et al., 2014). As we showed previously (Kubik et al., 2015), depletion of these PTFs alone results predominantly in increased average nucleosome occupancy in the vicinity of the PTF binding site, due either to encroachment of adjacent nucleosomes at an NDR or stabilization of a fragile nucleosome. Averaged nucleosome occupancy change +/- 500 bp from the binding sites of each PTF are shown in Figure 5A (see also Figure S5A). Importantly, double-depletion experiments did not reveal a fully epistatic relationship for any of the RSC-PTF combinations (Figure 5A), arguing against an obligatory role of a PTF in RSC recruitment, or vice versa.

**Figure 5.**
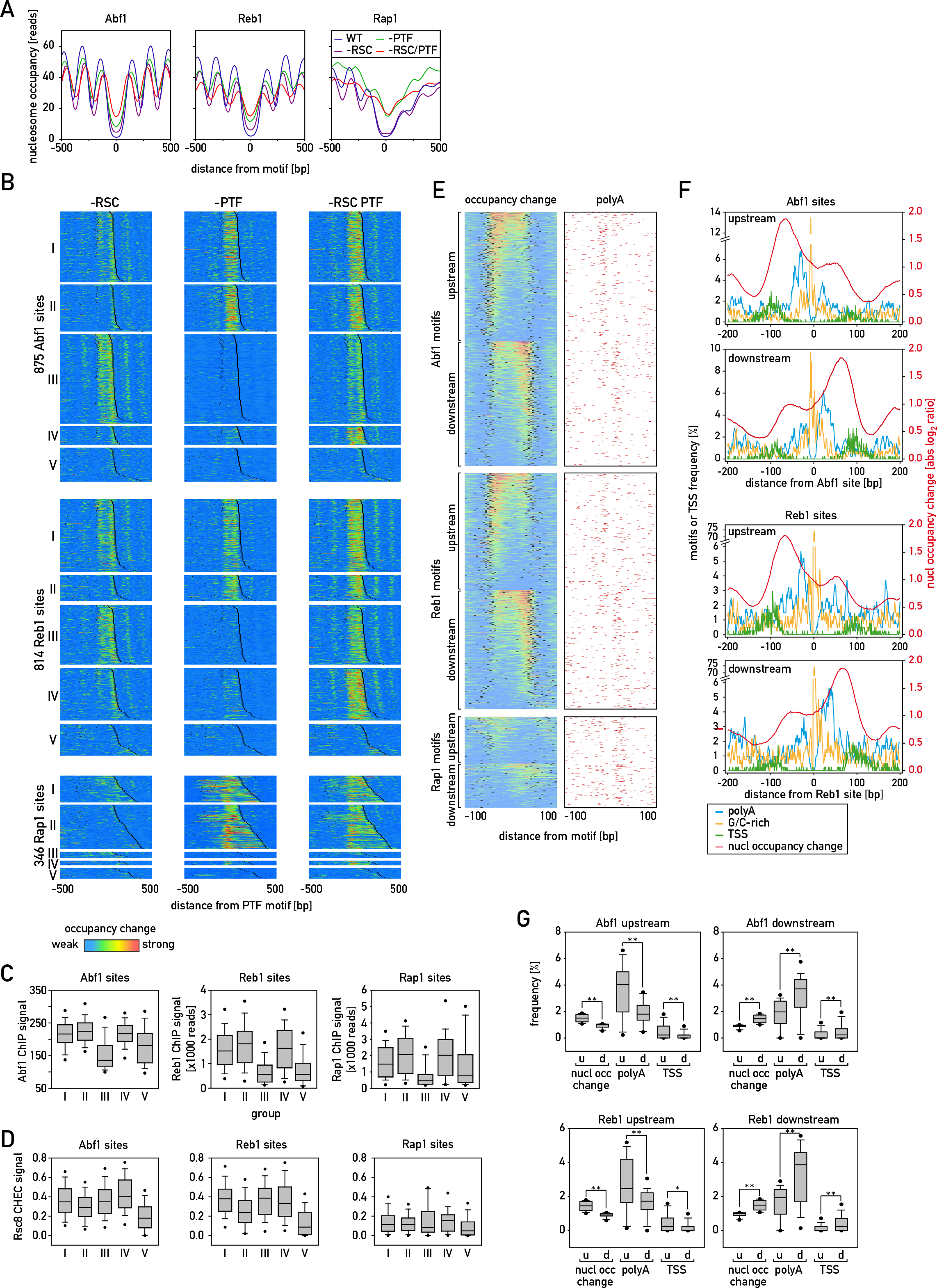
PTFs and RSC affect nucleosomes independently. (A) Average nucleosome occupancy plots, centered on PTF binding sites of Abf1 (left), Reb1 (middle) or Rap1 (right), in wild type cells (blue) and cells depleted of RSC (purple), respective PTF (green) or both RSC and PTF (red). (B) Heatmap showing absolute nucleosome occupancy change upon depletion of Sth1 (left), PTF (middle) or both factors simultaneously (right) in strains where the depleted PTF was: Abf1 (top), Reb1 (center) or Rap1 (bottom). Maps represent 1000 bp-wide regions centered on promoter PTF binding sites grouped according to the observed effects: I-additive effect of double depletion, II-no effect of RSC depletion, III-no effect of PTF depletion, IV-enhanced effect of double depletion, V-no effect of either depletion. The regions in each group were sorted according to the distance to the nearest TSS; TSS positions are represented by black boxes. (C) Boxplots showing ChIP-seq signal of PTFs in regions belonging to each of the groups described in (B) calculated -/+ 100 bp from Abf1, Reb1 or Rap1 motifs. (D) Boxplots showing normalized Rsc8 ChEC-seq signal of PTFs in regions belonging to each of the groups described in (B) calculated like in (C). (E) Heatmap showing absolute nucleosome occupancy change upon RSC depletion in 200 bp-wide regions centered on PTF binding sites (left) and polyA motif distribution in these regions (right); the regions were sorted by the strength of nucleosome occupancy change and divided based on whether the effect was stronger upstream or downstream from the PTF binding site; TSS positions are represented by black boxes. (F) Plots showing frequencies (per bp) of polyA motifs (blue), G/C-rich motifs (yellow), TSS (green) as well as average nucleosome occupancy change upon RSC depletion (red) in the regions bound by Abf1 or Reb1 shown in (E) where Sth1 depletion effects were more prominent either upstream or downstream from the motif. (G) Boxplots comparing the values for nucleosome occupancy change upon RSC depletion (calculated within 100 bp from PTF motif), polyA motif frequency per bp (within 50 bp) and TSS frequency per bp (within 150 bp) in the regions bound by Abf1 or Reb1 shown in (E).

By comparing nucleosome occupancy changes around each individual PTF motif upon depletion of the PTF and/or RSC we observed five types of behavior that were common to all three PTFs (Figure 5B; see Experimental procedures for details of this analysis). At many promoters both single depletions (PTF or RSC) yielded nucleosome rearrangements that were similar in magnitude and roughly additive in the double depletion experiment (group I), indicative of independent action, though not ruling out some degree of cooperativity between the PTFs and RSC. At PTF binding sites of groups II and III the nucleosome rearrangements were similar between double depletion and depletion of either the PTF or RSC alone, respectively, indicating that action of either the PTF or RSC was dominant and largely independent of the second factor. Interestingly, at a number of PTF binding sites the effects of depletion of both factors appeared stronger than the sum of single depletion effects (group IV) (see Figure S5A for some specific examples), suggesting that the factors act at least partially redundantly at these locations. Finally, a relatively small fraction of total sites displayed minor changes upon RSC depletion and little or no significant changes upon PTF depletion (group V).

To investigate what contributes to the different outcomes observed in each group we first focused on groups III and V, where the effects of single PTF depletion were minimal or non-existent. We noticed that the ChIP signal intensities (Kasinathan et al., 2014; Knight et al., 2014) in these two groups, which might reflect PTF binding strength and/or occupancy, were significantly lower than in the other groups (Figure 5C). This was true for all three PTFs. Consistent with these observations, motif comparisons revealed that sites in groups III and V had slightly lower information content in comparison to other groups (Figure S5B). Together with an overall strong correlation between ChIP signal intensity and nucleosome occupancy change upon PTF depletion (Figure S5C), these findings suggest that the weak effects of PTF depletion in groups III and V are a direct consequence of weaker PTF binding. Furthermore, we noted that for group III and V promoters the average signal of another, non-depleted PTF was often higher in these groups than in others (Figure S5D). Group IV sites for both Abf1 and Reb1 also showed relatively high signal for the other factor and a relatively small effect upon depletion of an individual PTF, suggestive of PTF redundancy at these promoters.

We observed a similar sequence-dependent binding difference for RSC that correlates with the strength of nucleosome occupancy changes measured in the different groups. Thus, the lack of a significant effect of RSC depletion at group V promoters, as well as those of group II for Abf1 and Reb1, is correlated with weaker RSC binding compared to other groups, as judged by Rsc8 ChEC signal intensity (Figure 5D). Consistent with this weaker ChEC signal we noted a paucity of polyA motifs surrounding both Abf1 and Reb1 sites in groups II and V, as well as fewer G/C-rich motifs at near Abf1 sites in these groups (Figure S5E,F). Taken together, this analysis suggests that PTF action is limited by binding site affinity and redundancy, whereas RSC action in the vicinity of some PTF sites is limited by motif frequency, in addition to the motif proximity effect described earlier (Figure 3).

We often observed that the effects of RSC depletion were stronger on one side of a PTF binding site than the other. To investigate what determines RSC activity towards particular nucleosomes in the vicinity of PTF binding sites we sorted the promoters bound by each PTF according to the magnitude of the nucleosome occupancy change following RSC depletion and then divided all promoters into two groups based upon whether the strongest effect was observed upstream or downstream of the PTF motif (assigning an arbitrary motif orientation; Figure 5E, left panel). When we then scored polyA tracts on both sides of the PTF motif it became clear that for both Abf1 and Reb1 sites, polyA tracts are more prevalent, on average, on the side of the PTF where RSC action is strongest (Figure 5F and G, right panel). G/C-rich motifs instead tended to map coincident with or very close to either side of PTF motifs (Figure 5F). In fact, Reb1 sites often contain a G/C-rich motif. Significantly, TSSs are more frequent at Abf1 and Reb1 flanks with polyA motifs, where they are positioned further downstream. Although both polyA and G/C-rich motifs are more frequent downstream from Rap1 sites, these regions are only weakly affected by RSC depletion, for reasons that are still unclear (Figure 5E and data not shown).

We also observed a distinct property of Rap1, which appears to act on nucleosomes in a directional manner, with little help from RSC, as pointed out above. This fact is seen when comparing Figure 5B (bottom panels), where the nucleosome maps surrounding the Rap1 motif have been oriented with respect to the nearest TSS (always to the right), and Figure S5G, where the regions are oriented with respect to the Rap1 motif itself. In the latter case, nucleosome occupancy displays prominent changes both at the Rap1 site itself and at one unique side of the motif, with only minor effects seen on the other side at a few promoters. These findings indicate that Rap1 is unique amongst the three PTFs tested in its ability to guide nucleosome rearrangements with a well-defined directionality, and may be related to the fact that Rap1 motifs at ribosomal protein gene (RPG) promoters predominantly display a unique orientation relative to the TSS (Knight et al., 2014).

### Chromatin changes provoked by PTF or RSC depletion correlate with reduced transcription

We first asked if PTFs, like RSC (see Figure 2), are implicated in regulating +1 nucleosome position and TBP binding. To do so, we measured both parameters following rapid nuclear depletion of each PTF alone or in combination with RSC. Indeed, following depletion of individual PTFs, we observed an upstream shift of a large fraction of the nearby +1 nucleosomes, which increased their proximity to, or overlap with, a TBS (Figure S6A). This effect was modestly exacerbated by simultaneous depletion of RSC (Figure S6A). To determine if PTF-mediated +1 positioning affects PIC assembly, we again performed TBP ChIP-seq experiments following depletion of individual PTFs or the PTF plus RSC. We observed a significant decrease in TBP binding at specific PTF target genes for all three PTF depletions, with or without simultaneous RSC depletion, that was strongly correlated with the change in nucleosome occupancy measured at the respective TBSs (Figure 6A). This correlation also held genome-wide (Figure S6B).

**Figure 6.**
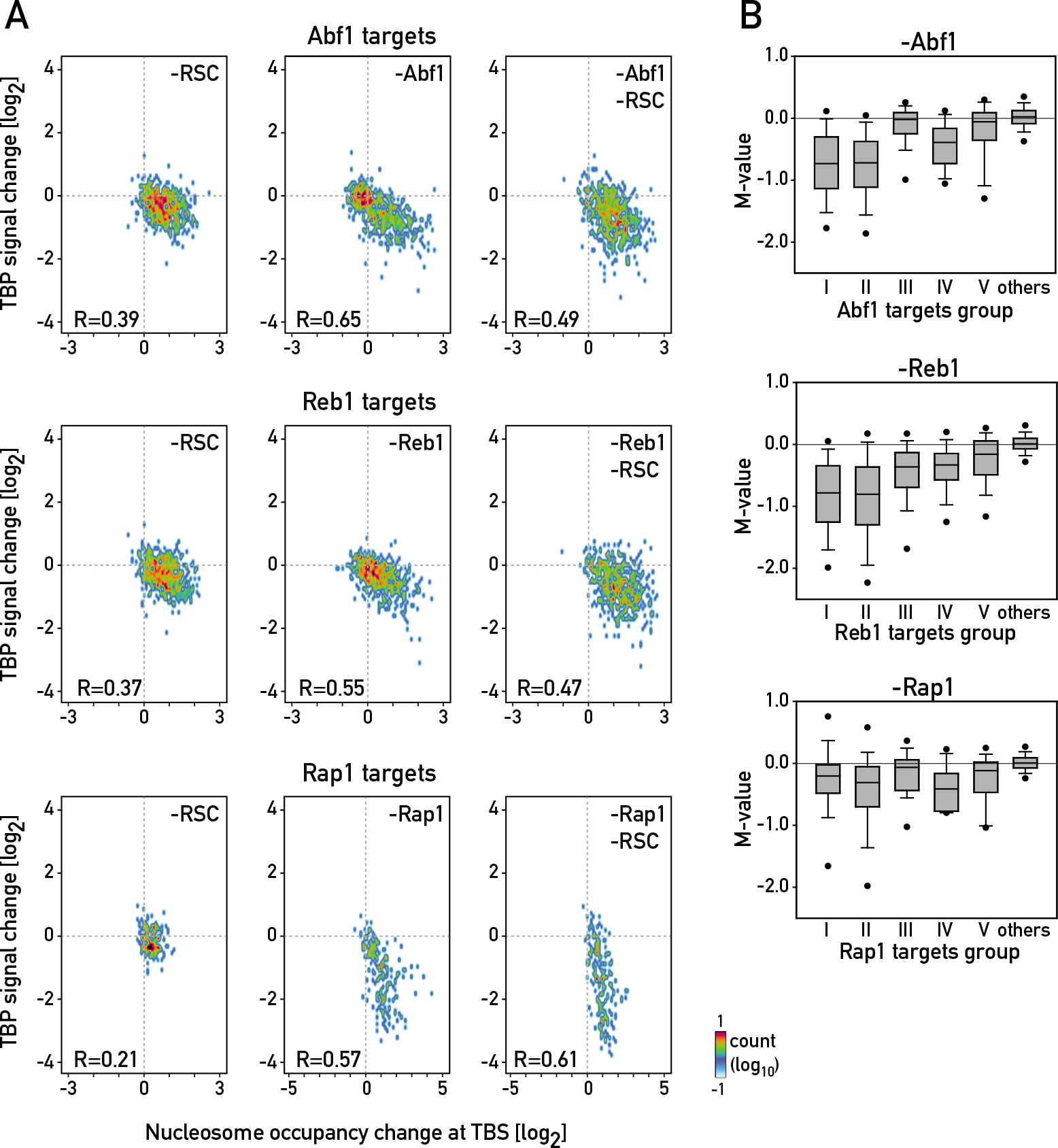
PTF and RSC-mediated organization of promoter nucleosomes is important for transcription. (A) Scatterplots showing the dependence of TBP signal change and nucleosome occupancy change upon depletion of RSC (left), PTF (middle) or each PTF in combination with RSC (right) calculated in 200 bp-wide regions surrounding TBSs of genes whose TSS was the closest to the PTF motifs. (B) Boxplots showing mRNA levels change of genes present in each motif group (I-V) shown in Figure 5B and all other genes (“others”), measured by microarray upon depletion of PTFs.

To extend the functional significance of these TBP binding changes, we measured gene expression following depletion of each PTF. Depletion of Abf1, Reb1 or Rap1 led to significant decreases in the mRNA levels of the corresponding target genes in groups I, II and IV (Figure 6B). In contrast, target genes of groups III and V, where nucleosome occupancy and TBP binding were essentially unaffected by PTF depletion, showed little or no transcriptional down-regulation. Moreover, genes whose promoters were not bound by any of the tested PTFs did not display transcriptional changes (Figure 6B, “others”). An independent measurement of transcription in RSC-depleted cells (RNAP II ChIP-seq analysis) also bore out the correlation between nucleosome occupancy changes and transcription (Figure S6C). Thus group III genes generally showed the strongest decrease in RNAP II levels following RSC depletion, while group II genes bearing Abf1 and Reb1 motifs were the least affected. Taken together, these data strongly support the notion that the PTFs and RSC act through +1 nucleosome positioning to promote TBP binding, thus stimulating PIC assembly and transcription initiation.

## Discussion

### Promoter nucleosome positioning is intimately connected to TBP binding and transcription

PIC formation is a key early event in transcription initiation by RNAP II. In vitro studies indicate that initiation of transcription is inhibited when a nucleosome is placed precisely on the TATA-box and TSS (Lorch et al., 1987), probably due to reduced TBP binding (Godde et al., 1995). These observations have led to the hypothesis that nucleosome-regulated TATA box accessibility plays an important role in gene activation (Lomvardas and Thanos, 2001; Struhl, 1999), particularly for a subset of highly inducible or stress-related genes bearing a canonical TATA box in their promoter (Basehoar et al., 2004; Tirosh and Barkai, 2008; Zaugg and Luscombe, 2012). The more recent finding that TBP also binds to the TATA-like elements often found at highly expressed TFIID-dominated housekeeping genes (Rhee and Pugh, 2012) suggests that nucleosome positioning might affect PIC assembly more globally. However, these ideas have been difficult to test in vivo, in large part due to the fact that many of the proteins implicated in promoter nucleosome positioning, such as PTFs, RSC and TBP, are essential for cell viability.

We show here that both RSC and a group of three PTFs contribute to positioning of +1 nucleosomes at a large number of RNAP II promoters in yeast, such that their activity displaces the TBS further away from the dyad axis of the +1 nucleosome, in many cases shifting the TBS into a nucleosome-free region. This effect is strongly correlated with increased TBP binding and transcription at more than half of all protein-coding genes in cells growing under optimal conditions. We propose that the effect of both RSC and PTFs on +1 nucleosome positioning is direct and should be considered as an early step in the promotion of PIC assembly and subsequent RNAP II recruitment (summarized graphically in Figure 7). In support of this notion, we note that rapid nuclear depletion of RNAP II itself does not lead to any upstream shift of +1 nucleosomes genome-wide (Kubik et al., 2015; Weiner et al., 2010), arguing against the notion that +1 displacement is a consequence of, rather than a cause of, transcription initiation. Similarly, TBP binding itself does not influence +1 nucleosome position at the vast majority of genes.

**Figure 7.**
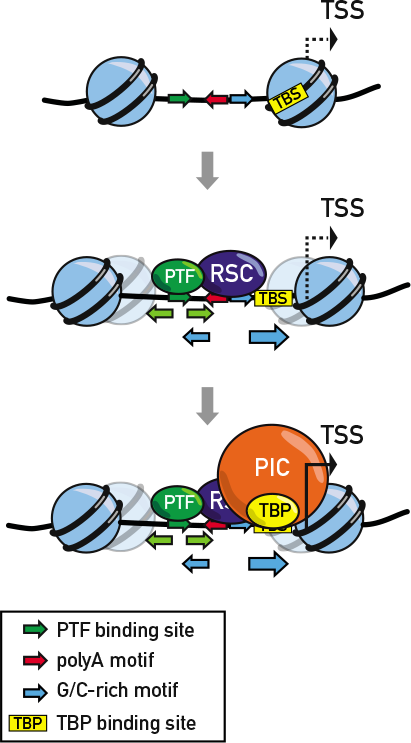
Arrangement of promoter motifs determines nucleosome architecture and drives transcription. Model showing how the arrangement of PTF, G/C-rich and polyA motifs (top) largely determines nucleosome positions at the promoter by driving the binding and activity of PTFs and RSC (middle). The orientation of polyA directs RSC-mediated remodeling in the 5’ direction of the motif. Activity of both PTFs and RSC independently leads to exposition of the TBP binding site and PIC formation (bottom), which facilitates transcription.

Our findings are consistent with previous reports in which ts-lethal mutants were used to inactivate either Rap1 or Abf1, which was shown to lead to both chromatin changes and transcriptional down-regulation at the respective target genes (Badis et al., 2008; Ganapathi et al., 2011; Yarragudi et al., 2004), although in these studies the mechanistic link to transcription was unclear. The importance of the +1 nucleosome shift in gene activation that we observe here is also consistent with the recent finding of pervasive regulation of +1 positions in cells passaging through the yeast metabolic cycle, where periodic expression of a large fraction of the genome is observed (Nocetti and Whitehouse, 2016). Since these studies did not involve genetic perturbations, it was difficult to infer underlying mechanisms.

### The role of DNA sequence in shaping promoter nucleosome architecture

We showed previously that the nucleosome landscape upstream of the TSS-associated +1 nucleosome at a large fraction of all RNAP II-transcribed genes in yeast, whether nucleosome-depleted or occupied by an MNase-sensitive FN, is strongly influenced by the essential RSC remodeler. Our data suggested that RSC action in vivo is promoted by two short DNA sequence motifs, and serves to “push” the +1 and −1 nucleosomes away from each other (Kubik et al., 2015). Here we present more detailed evidence indicating that the polyA and G/C-rich motifs driving RSC binding and/or action, when in close proximity (<40 bp apart), operate together and lead to strong nucleosome displacement. We find that RSC acts more strongly overall when the G/C-rich motif is on the 5’ side of the polyA motif. Related to this observation, we also show that nucleosome displacement is asymmetric with respect to the motifs, with a stronger effect observed on the 5’ side of polyA motifs, recapitulating a recent observation made by studies of an in vitro nucleosome assembly system using purified components, including RSC and various PTFs (Krietenstein et al., 2016). Our findings may explain the earlier observation of an asymmetric arrangement of nucleosomes around polyA tracts in vivo (de Boer and Hughes, 2014; Wu and Li, 2010). Interestingly, recent work shows that polyA affects the direction of nucleosome sliding by the chromatin remodeler Chd1, which belongs to a subfamily distinct from RSC both functionally and mechanistically (Winger and Bowman, 2017). Overall, these observations suggest that DNA sequence might, apart from affecting nucleosome positioning through direct effects related to helix bending or specific DNA-histone interactions (Struhl and Segal, 2013) also influence nucleosome positioning through pervasive effects on remodeler binding and activity.

Although our conclusions regarding the effect of motif spacing and orientation on RSC action are based upon correlation, and warrant testing by more systematic analyses connecting sequence to function, they are strongly buttressed by our measurements of Rsc8 binding using the ChEC assay. Indeed, we show that this novel approach to localize and quantify RSC binding in vivo correlates extremely well with motif arrangement and strength of RSC action, as inferred by rapid depletion experiments. By contrast, native ChIP-seq of Sth1 yields a weaker correlation with RSC action, and standard ChIP-seq of Rsc8 is considerably worse than either alternative method (Figure S3B). Furthermore, our findings indicate that ChEC may also reveal more fine-structure details of RSC binding (Figure 4A).

### Functional interactions of pioneer factors and RSC remodeler

Our data reveal a highly stereotypical arrangement of polyA and G/C-rich motifs surrounding two of the more ubiquitous and abundant PTFs, Abf1 and Reb1. G/C-rich motifs cluster very close to both Abf1 and Reb1 binding sites, whereas polyA tracts are more distant, typically at 20-80 bp from the PTF. Notably, there is a clear tendency for RSC to more strongly alter nucleosome occupancy on the side of the PTF containing a polyA tract, where the TSS is located. The conserved nature of this arrangement is underscored by the observation that both Abf1 and Reb1 sites associated with divergent transcription often contain polyA tracts on both sides of the PTF site and strong bilateral nucleosome occupancy changes.

By comparing effects of RSC or PTF nuclear depletion alone to that of double depletions, we begin to obtain some insight into the functional relationship between these two classes of chromatin-modulating factors. One finding that emerges from these experiments is that neither factor would appear to be completely dependent upon the other for its ability to provoke nucleosome occupancy changes. Thus, in the cases where depletion of RSC has little additional effect over PTF depletion the promoter in question often lacks closely spaced polyA and G/C-rich motifs and is only weakly, if at all, affected by RSC depletion alone. We conclude that RSC is not dependent upon a PTF for promoter binding, either through a recruitment mechanism or by clearing nucleosomes from polyA or G/C-rich motifs. Conversely, cases where Abf1, Reb1 or Rap1 depletion causes little or no additive effect with RSC are often associated either with weak binding or the presence of another PTF nearby. Thus, none of the three PTFs tested appear to require RSC to expose its binding site within chromatin, hence justifying their classification as PTFs. Furthermore, these findings suggest that PTFs can act redundantly at some promoters, though this awaits a direct test. Nevertheless, even though no factor is epistatic to another, the data do not rule out the possibility that each helps to promote the action of the other, at least to some extent.

The additive (group I) or synergistic (group IV) effects of simultaneous PTF and RSC depletion suggest that these factors possess at least partially independent functions, consistent with the fact that the former operate through tight binding to a specific recognition sequence whereas the latter can engulf a nucleosome and displace it directly through the use of an ATP-dependent motor (Clapier et al., 2017). One attractive model to explain the independent but coordinated action of PTFs and RSC would be that PTFs, though strong DNA binding, generate a barrier to nucleosome binding or encroachment that RSC might utilize to drive nucleosomes away from the site of PTF binding. Such a model might be testable in vitro.

Finally, we note that Rap1, unlike Abf1 and Reb1, guides nucleosome rearrangements with a well-defined directionality, consistent with the fact that Rap1 motifs at ribosomal protein gene (RPG) promoters are predominantly well oriented towards the TSS of these genes (Knight et al., 2014). Strikingly, RPG promoters whose TSSs are located 300 to 500 bp downstream from Rap1 binding sites (category I promoters; (Knight et al., 2014)) display prominent nucleosome occupancy changes upon depletion of this PTF (Kubik et al., 2015). Since it is unlikely that Rap1, lacking ATPase activity, can directly act as far as 500 bp from its binding site, we suggest that Rap1 might recruit multiple chromatin remodelers at category I genes to destabilize or displace the +1 nucleosome. The contribution of other factors to nucleosome positioning at RPG promoters seems relatively minor. For example, high mobility group protein Hmo1 binds to G/C-rich regions at category I promoters but its role is limited to fine tuning +1 nucleosome position (Kasahara et al., 2011; Reja et al., 2015), since its deletion produces only a small +1 shift, comparable to that of TBP depletion, but considerably less than that caused by removal of Rap1.

### Additional factors determining +1 nucleosome position

Although our data demonstrate a pervasive role for both PTFs and RSC in downstream displacement of the +1 nucleosome to a position more favorable for TBP binding, they do not rule out a role for other factors. Indeed, two other nucleosome remodelers, ISW1 and ISW2, have been implicated in upstream movement of the +1 nucleosome (Parnell et al., 2015; van Bakel et al., 2013; Whitehouse et al., 2007; Yen et al., 2012), and may thus counteract PTF/RSC action at many promoters. Although neither of these remodelers is essential for viability, unlike the PTFs and RSC, their action may be highly redundant. It will be interesting to test these ideas by multiple depletion experiments followed by +1 nucleosome mapping. TBP is another factor that has been proposed to influence +1 nucleosome positioning. However, our anchoring experiments indicate that TBP plays a role in +1 positioning at only a limited set of genes dominated by those encoding ribosomal proteins. We speculate that this may be linked to the observation that Rap1, which serves as the PTF at many of these genes, can directly interact with both TBP and several TAF proteins in TFIID (Bendjennat and Weil, 2008; Garbett et al., 2007). It is also possible that the +1 nucleosome simply moves to a thermodynamically favorable position determined by local DNA sequence features following removal of both RSC and an upstream PTF. In this scenario, one could imagine that other remodelers such as ISW1 or ISW2 act only to stabilize a default +1 position when PTF and RSC are removed. Another layer of regulation that needs to be considered is related to the fact that a well positioned +1 nucleosome poses a barrier for initiating polymerase which needs to be bypassed (Weber et al., 2014).

### Is +1 positioning by PTFs and RSC part of a global gene regulatory mechanism?

Our results indicate that transcription initiation is stimulated genome-wide by the combined action of RSC and a relatively small number of PTFs. More specifically, our findings suggest that the cell consumes ATP, through the RSC remodeler, to actively displace the +1 nucleosome to a position more favorable for TPB binding. The extent to which this occurs in coordination with, or opposition by other remodelers, is still unclear. In any event, this begs the question of why, under optimal growth conditions, an energy-consuming process is required to establish or maintain a chromatin configuration that supports wild-type levels of transcription. One interesting possibility is that this mechanism provides a direct link between the metabolic state of the cell and genome-wide transcription initiation rates. A global mechanism for growth control of gene activation through promoter nucleosome positioning has recently been suggested from studies of yeast cells entrained to an ultradian metabolic cycle (Nocetti and Whitehouse, 2016). It remains to be determined to what extent the recruitment of PTFs and RSC itself is regulated rather than constant throughout various physiological conditions. The functional studies described here implicate both RSC remodeler and pioneer transcription factors in such a mechanism and point to future lines of research aimed at better understanding how genome-wide control of promoter nucleosome architecture is linked to cell growth.

## Author contributions

Conceptualization, S.K. and D.S.; Formal Analysis, S.K., E.O. W.J. and J.F.; Investigation, S.K., E.O., S.M., B.A. and M.J.B.; Data Curation, S.K and W.J.; Writing – Original Draft, S.K. and D.S.; Funding Acquisition, D.S. and F.C.P.H.; Resources, D.S. and F.C.P.H.; Supervision, D.S. and F.C.P.H.

## Acknowledgements

We would like to thank Mylène Docquier and the iGE3 Genomics Platform of the University of Geneva for high throughput sequencing services, Nicolas Roggli for expert assistance with data presentation and artwork, Florian Steiner for comments on the manuscript, and all members of the Shore lab for comments and discussions throughout the course of this work. M. J. B. was supported in part by an iGE3 Ph.D. student fellowship. D. S. would like to acknowledge funding for this work from the Swiss National Science Foundation and the Republic and Canton of Geneva.

## STAR Methods

### CONTACT FOR REAGENT AND RESOURCE SHARING

Further information and requests for resources and reagents should be directed to and will be fulfilled by the Lead Contact, David Shore.

### METHOD DETAILS

#### Yeast growth conditions

Yeast strains used in this study are listed in Key Resources Table. Strains were grown in YPAD at 30°C. Anchor-away of FRB-tagged protein was induced by the addition of rapamycin (1 mg/ml of 90% ethanol/10% Tween stock solution) to a final concentration of 1 µg/ml for 1h (Haruki et al., 2008).

#### MNase-seq

MNase-seq experiments were performed essentially as described previously (Kubik et al., 2015). Yeast cultures were grown at 30°C in YPAD to OD_600_~0.4 and crosslinked with 1% formaldehyde for 5 min. at room temperature. Cell pellets were resuspended in spheroplasting buffer (1M sorbitol, 1 mM β-mercaptoethanol, 10 mg/ml zymolyase; 1 ml of buffer per 100 ml of cell culture) and incubated for 8 min. at room temperature. Spheroplasts derived from the cultures were washed twice with 1 ml of 1 M sorbitol and treated with different concentrations of MNase, ranging from 0.1 to 3 units. One concentration of MNase was applied to spheroplasts derived from an equivalent of 25 ml of culture. The samples were incubated at 37°C for 45 minutes. The reactions were stopped by addition of EDTA (30 mM final conc.) and the crosslinks were reversed by addition of SDS (0.5% final conc.) and proteinase K (0.5 mg/ml final conc.), incubation at 37°C for 1h followed by incubation at 65°C for at least 2h. Samples underwent phenol-chloroform extraction and DNA was ethanol precipitated from the aqueous phase. After washing with 70% ethanol the DNA pellet was dried, resuspended in 30 µl TE with RNase and the samples were incubated for 30 min. at 37°C to digest RNA. 2 µl aliquots were resolved on 3% agarose gel to evaluate the digestion. It was critical at this step that the samples from anchor away experiments treated with vehicle and rapamycin that were to be compared were digested to exactly the same extent, as evaluated by the intensity ratios between mono-and di-nucleosomal DNA bands.

DNA from MNase-digested chromatin was prepared for deep sequencing according to (Henikoff et al., 2011). First, DNA was incubated with an end-repair mixture containing 0.4 mM dNTPs, 7.5 U T4 DNA Polymerase, 2.5 U of Klenow fragment and 25 U of T4 PNK for 30 min. at 20°C (reaction volume 50 µl). Then, DNA was extracted with phenol-chloroform and purified using Illustra MicroSpin S-300 HR columns (GE Healthcare). A overhangs were added to DNA fragments by incubation with 0.2 mM dATP and 15 U of Klenow exo-for 15 min. at 37°C. DNA was extracted and purified using MicroSpin columns again. The TruSeq adaptors were ligated using 3000 units of Rapid Ligase (Enzymatics) for 15 min. at 20°C. DNA was purified using AMPure XL beads (Agencourt) (1:1 ratio of beads to DNA prep) according to manufacturer’s description. The libraries were amplified using KAPA Polymerase (17 cycles of 98°C 20s/60°C 15s/72°C 15s) and purified again using the AMPure XL beads (1:1 ratio).

The libraries were sequenced using HiSeq 2500 in the paired-end mode. Mapping of the sequencing data to sacCer3 genome assembly was performed using HTSStation (David et al., 2014). Mapped reads were trimmed by 15 bp from each side to better visualize individual nucleosome peaks.

#### ChIP-seq

Cultures of 240 ml in YPAD were collected at OD_600_ 0.5-0.8 for each condition. The cells were crosslink ed with 1% formaldehyde for 5 min. at room temp. and quenched by adding 125 mM glycine for 5 min. at room temp. Cells were washed with ice-cold HBS and resuspended in 3.6 ml of ChIP lysis buffer (50 mM HEPES-Na pH 7.5, 140 mM NaCl, 1mM EDTA, 1% NP-40, 0.1% sodium deoxycholate) supplemented with 1mM PMSF and 1x protease inhibitor cocktail (Roche). Samples were aliquoted in 6 eppendorf tubes and frozen. After thawing the cells were broken using Zirconia/Silica beads (BioSpec). The lysate was spun 13′000 rpm for 30 min. at 4°C. The pellet was resuspended in 300 µl ChIP lysis buffer + 1mM PMSF and sonicated for 15 min. (30” On - 60” Off) in the Bioruptor (Diagenode). The lysate was spun at 7000 rpm for 15 min. at 4°C. The antibody (1 µg / 300 µl of lysate) was added to the supernatant and incubated for 1h at 4°C. The magnetic beads were washed three times with PBS plus 0.5% BSA and added to the lysates (30 µl of beads/300 µl of lysate). The samples were incubated for 2h at 4°C. The beads were washed twice with (50 mM HEPES-Na pH 7.5, 140 mM NaCl, 1mM EDTA, 0.03% SDS), once with AT2 buffer (50 mM HEPES-Na pH 7.5, 1 M NaCl, 1mM EDTA), once with AT3 buffer (20 mM Tris-Cl pH 7.5, 250 mM LiCl, 1mM EDTA, 0.5% NP-40, 0.5% sodium deoxycholate) and twice with TE. The chromatin was eluted from the beads by resuspending them in TE+1% SDS and incubation at 65°C for 10 min. The eluate was transferred to an Eppendorf tube and incubated overnight at 65 °C to reverse the crosslinks. The DNA was purified using High Pure PCR Cleanup Micro Kit (Roche). The libraries were prepared using TruSeq ChIP Sample Preparation Kit (Illumina) according to manufacturer’s instructions. The libraries were sequenced using HiSeq 2500 and the reads were mapped to sacCer3 genome assembly using HTSStation (shift=150 bp, extension=50 bp).

#### ChEC-seq

The experiments were performed essentially as described in Zentner et al., (2015) with the following modifications. Cells in which MNase was fused at the C-terminus of Rsc8 were used to determine binding sites of the RSC remodeler. Cells in which MNase was placed under the control of *REB1* promoter were used as a control. One sample corresponds to 12 ml of culture at OD600=0.7. Cells were washed twice with buffer A (15 mM Tris 7.5, 80 mM KCl, 0.1 mM EGTA, 0.2 mM spermin, 0.5 mM spermidine, 1xRoche EDTA-free mini protease inhibitors, 1 mM PMSF) and resuspended in 200 µl of buffer A with 0.1% digitonin. The cells were incubated for 5 min at 30°C. Then, MNase action was induced by addition of 2 mM CaCl2 and stopped at desired timepoint by adding EGTA to a final concentration of 50 mM. DNA was purified using MasterPure Yeast DNA purification Kit (Epicentre) according to manufacturer’s instruction. Large DNA fragments were removed by a 5-minute incubation of DNA preps with 2.5x volume of AMPure beads (Agencourt) after which the supernatant was kept and MNase-digested DNA was precipitated using isopropanol. Libraries were prepared using NEBNext kit (New England Biolabs) according to manufacturer’s instructions. Before the PCR amplification of the libraries small DNA fragments were selected by a 5 minute incubation with 0.9x volume of the AMPure beads after which the supernatant was kept and incubated with the same volume of beads as before for another 5 min. After washing the beads with 80% ethanol the DNA was eluted with 0.1x TE and PCR was performed. Adaptor dimers were removed by a 5 minute incubation with 0.8x volume of the AMPure beads after which the supernatant was kept and incubated with 0.3x volume of the beads. The beads were washed twice with 80% ethanol and DNA was eluted using 0.1x TE. The quality of the libraries was verified by running an aliquot on a 2% agarose gel. Libraries were sequenced using HiSeq 2500 in single-end mode. Read ends were considered to be MNase cuts and were mapped to the genome (sacCer3 assembly) using HTSStation.

#### mRNA measurements

Gene expression upon PTFs depletion was measured by microarray analysis as described before (Kemmeren et al., 2014).

### QUANTIFICATION AND STATISTICAL ANALYSIS

#### Nucleosome analysis

TSS positions were taken from van Bakel et al (2013) and corrected for the genes with leader introns (taken from the UCSC database). Nucleosome dyad positions were determined using Peak Locator (minimum distance 100 bp, minimum signal=8 reads/10M) as described previously (Kubik et al., 2015). +1 nucleosome was defined as the dyad closest to TSS. Shifts of +1 nucleosomes were calculated by finding, in the rapamycin treated samples, the positions of nucleosomes closest to positions of +1 found in mock-treated cells. Manual inspection of nucleosomes where calculated shift upon Sth1 or PTF depletion was >30 bp in the downstream direction revealed that it was the +2 nucleosome position that was found. Therefore, for these positions we considered the closest nucleosome upstream from wild-type +1 dyad position to be the shifted +1. +1 nucleosome shifts were considered to be significant in a range between 25 and 105-120 bp, depending on the experiment, with smaller shifts considered as irrelevant (low confidence) and larger as false-positives. Nucleosome occupancy change was calculated as log2 ratio between nucleosome occupancy in cells depleted of a factor and mock treated cells. Absolute value of the log2 ratio was used where indicated to account for simultaneous occupancy increase and decrease at adjacent regions following nucleosome shift.

#### TBP binding

TBS positions were taken from (Rhee et al., 2012). When plotting TBS distribution (Figures 1C and S6A) we excluded genes where +1 nucleosome is found >200 bp from TSS. TBP occupancy was calculated in 200 bp-wide regions centered on TBSs.

#### ChEC-seq analysis

Cuts caused by free MNase displayed an increase in intensity in accessible regions of the genome (i.e. promoters) throughout the time-course, indicating that MNase cleaves in accessible regions with a probability that is proportional to the reaction time. In contrast, cleavage in the Rsc8-MNase strain appeared to decrease with increasing treatment time. We interpret this as an indication that Rsc8-MNase cuts rapidly at the promoters where RSC is specifically bound, leading to extensive degradation and consequent disappearance of signal after prolonged treatment. Therefore, as a measure of Rsc8 binding we used the signal obtained for Rsc8 after 10s of CaCl_2_ treatment normalized by dividing it by the signal obtained for free MNase after 20 min of CaCl_2_ treatment. For the heatmaps representing ChEC signal in Figures 3A (middle panel), 4A and 4B, when calculating signal ratio we excluded regions where the signal was lower than 10 reads/10M in order to decrease the background noise.

#### PTF motif analysis

The PTF binding motifs were taken from (Kasinathan et al. 2014) for Abf1 and Reb1 and from (Knight., et al 2014) for Rap1. Only motifs found at most 400 bp from TSS were considered. Absolute nucleosome occupancy change upon depletion of Sth1, a PTF or both factors simultaneously was calculated in 200 bp-wide regions centered on the motifs. Since Rap1 displayed nucleosome changes further from its binding site than Abf1 and Reb1 the region considered was 400 bp wide for this factor. If the average occupancy change was higher than 1 upon depletion of both – Sth1 and PTF or in the double depletion the motif was assigned to group I, if the change was >1 for either PTF or Sth1 depletion only, the motif was assigned to group II or III, respectively. Furthermore, if the ratio was higher upon double depletion than the sum of ratios of single depletions the motif was ascribed to group IV. The remaining motifs (group V) did not display any significant occupancy changes in our experiments. Motifs in each group were sorted according to their distance to the closest TSS unless stated otherwise.

R-values for all correlation measurements are Pearson’s correlation coefficients. Statistical significance of differences between groups of parameters was evaluated using Mann-Whitney Rank Sum test. Some of the plots were prepared using EaSeq (Lerdrup et al., 2016).

### DATA AND SOFTWARE AVAILABILITY

All sequencing and microarray data generated in this study was submitted to the GEO database as SuperSeries GSE98260.

**Figure S1.**
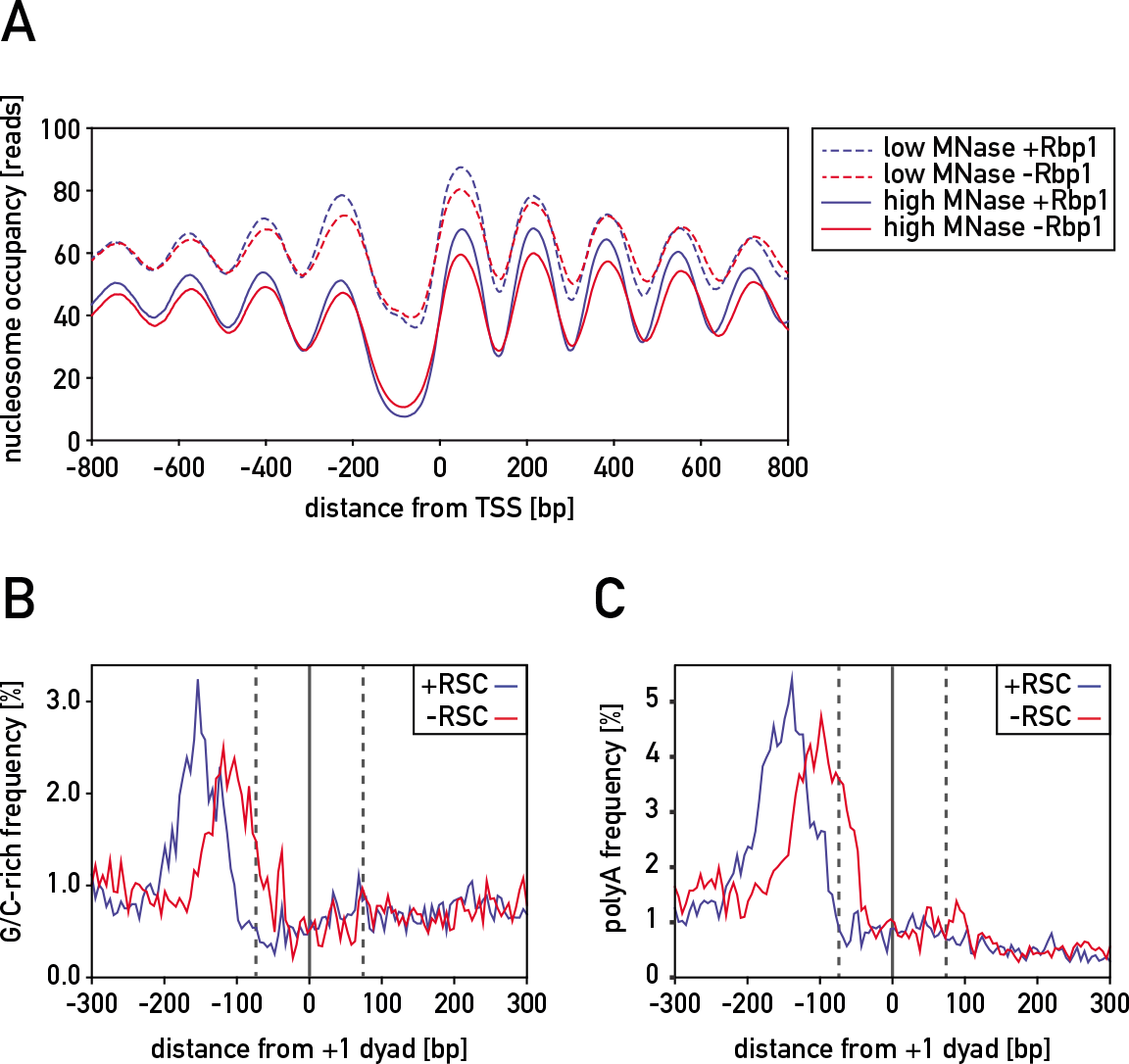
Related to Figure 1. **(A)** Average nucleosome occupancy obtained with either low (dashed lines) or high (solid lines) MNase concentration in the presence (blue) or absence (red) of RNAPII catalytic subunit Rbp1 centered on TSS of 1627 genes displaying, after Sth1 depletion, an increase in promoter (region spanning 400 bp upstream from TSS) nucleosome occupancy of at least 5 reads/bp. **(B)** Frequency of G/C-rich motif at genes whose +1 nucleosome significantly changed position after depletion of Sth1 in the presence (blue) and absence (red) of RSC, centered on +1 dyad in either condition. **(C)** Frequency of polyA motif presented as in (B).

**Figure S2.**
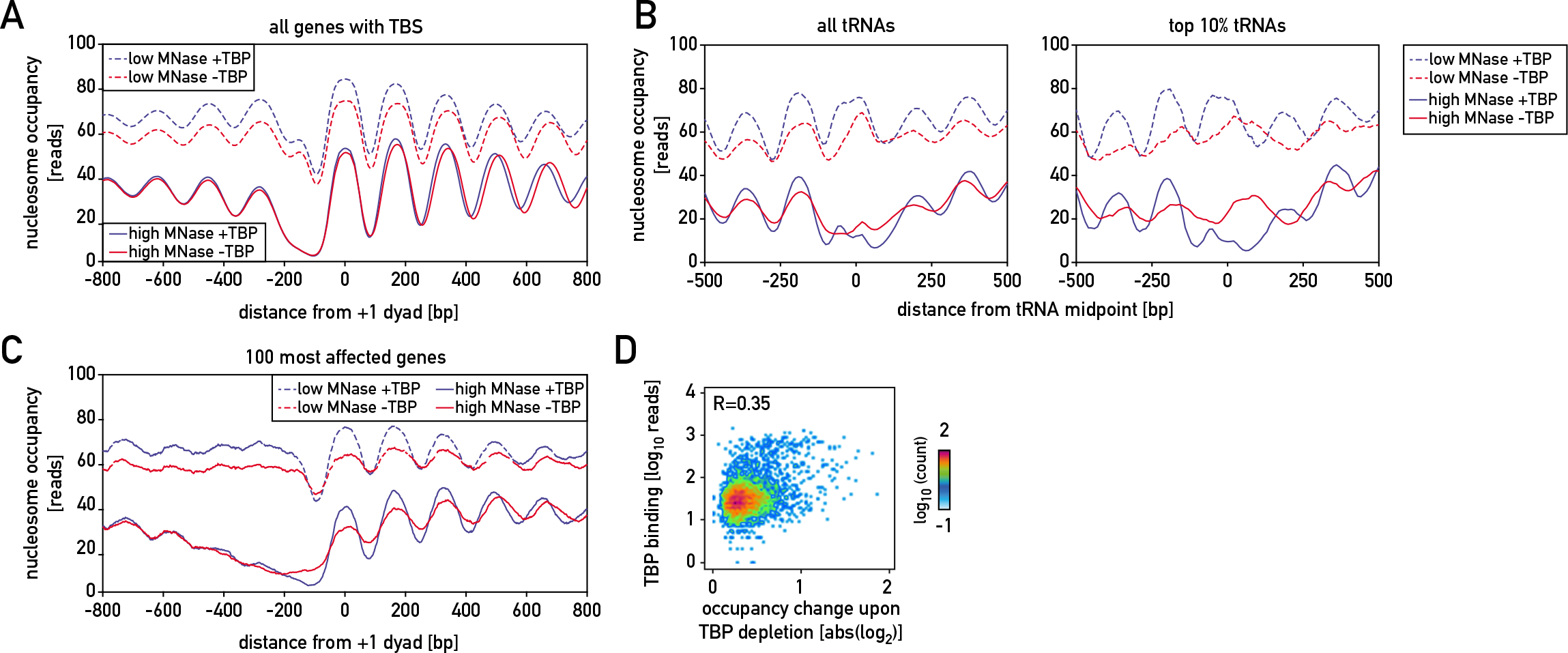
Related to Figure 2. **(A)** Average nucleosome occupancy plots in the wild-type cells (blue) and cells depleted of TBP (red) obtained with low (dashed lines) or high (solid lines) MNase concentration centered on +1 nucleosome dyad of all genes with an identified TBS (left). **(B)** Same as (A) but for all (left) or 10% most affected (right) tRNA genes. **(C)** Same as (A) but for 100 most affected genes, as determined by nucleosome occupancy change in the region spanning 200 bp upstream from +1 nucleosome dyad. **(D)** Scatterplot showing relationship between nucleosome occupancy change upon TBP depletion and TBP binding calculated for 200 bp-wide regions centered on TBS.

**Figure S3.**
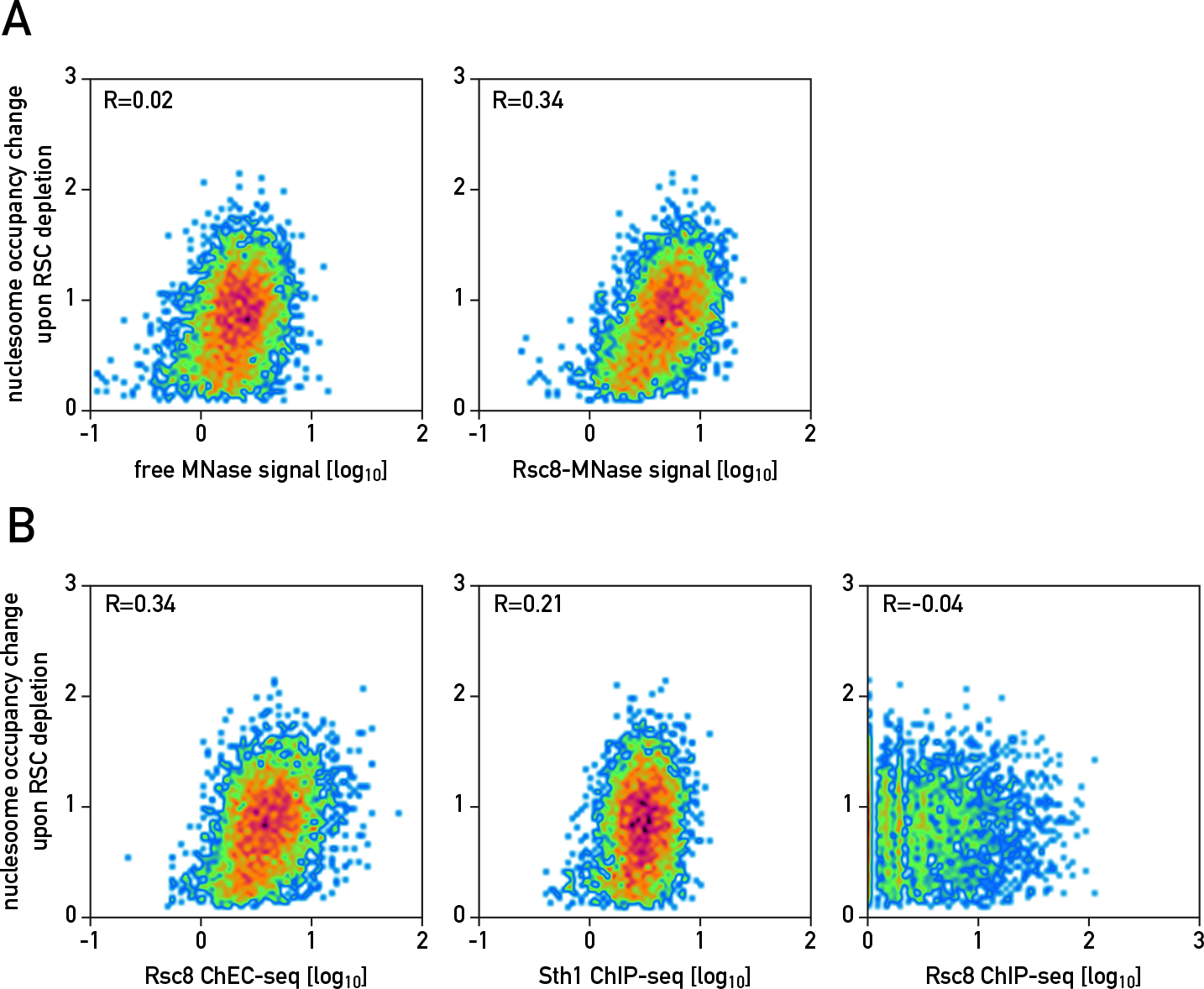
Related to Figure 3. **(A)** Scatterplots showing the relationship between free MNase (left) or Rsc8-MNase (right) ChEC signal and absolute nucleosome occupancy change upon RSC depletion calculated in regions from −200-0 bp with respect to +1 nucleosome dyad. **(B)** Scatterplots showing the relationship between nucleosome occupancy change upon RSC depletion and Rsc8 ChEC signal normalized to the free MNase signal (left), Sth1 native ChIP-seq signal (Ramachandran et al., 2015) (middle) and Rsc8 MNase ChIP-seq signal (Yen et al., 2012) (right).

**Figure S4.**
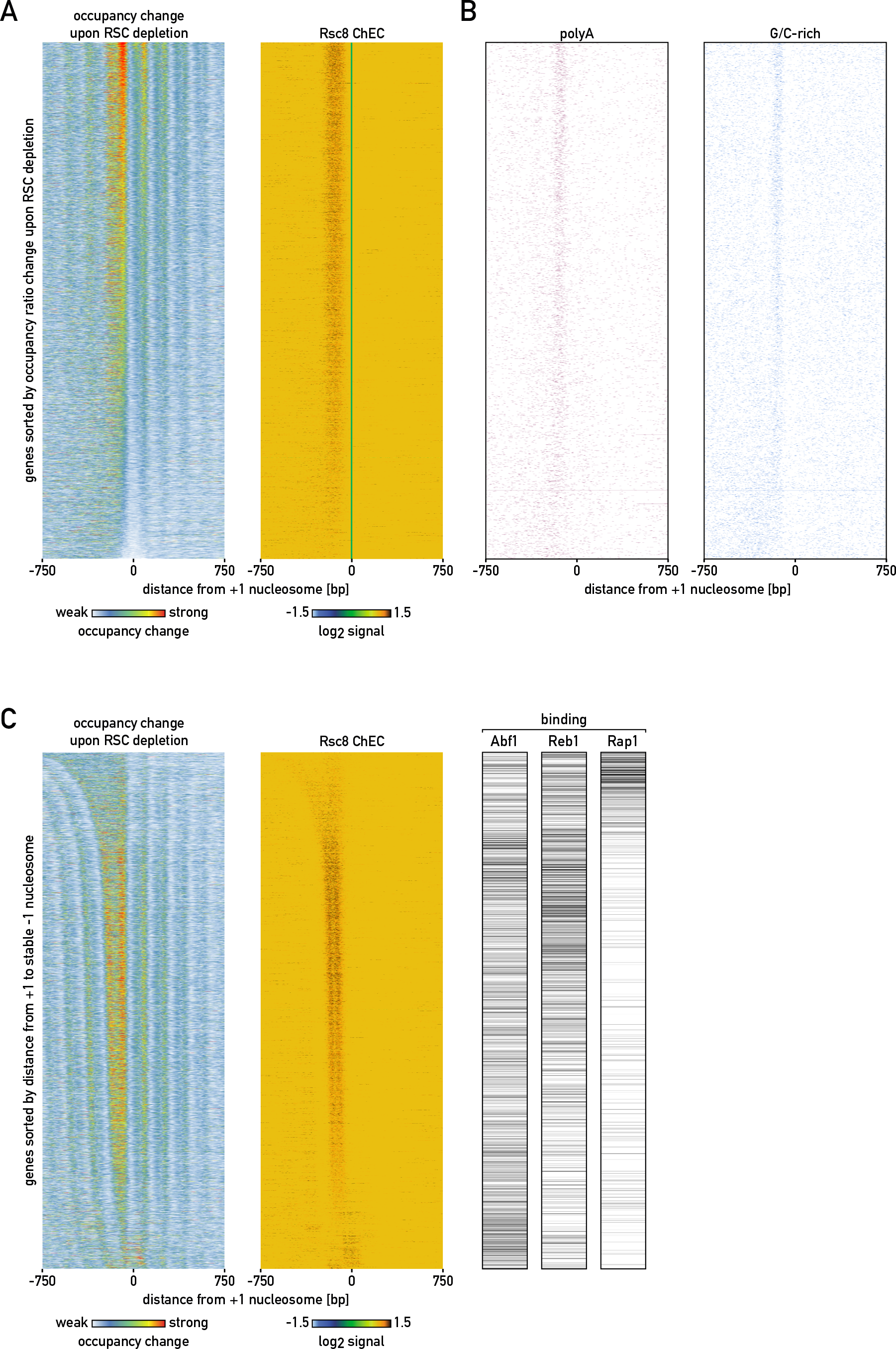
Related to Figure 4. **(A)** Heatmaps showing nucleosome occupancy change upon RSC depletion (left) and Rsc8 ChEC signal (normalized to free MNase) (right) in regions centered on +1 nucleosome of RNAPII genes, maps were sorted by the average nucleosome occupancy change in the region 200 bp upstream from +1 dyad. **(B)** Map showing distribution of polyA and G/C-rich motifs in the regions shown in (A). **(C)** (left and middle) Heatmaps as in (A) but sorted by the distance from +1 to first upstream stable nucleosome; (right) binding signal for 3 PTFs measured in the promoter.

**Figure S5.**
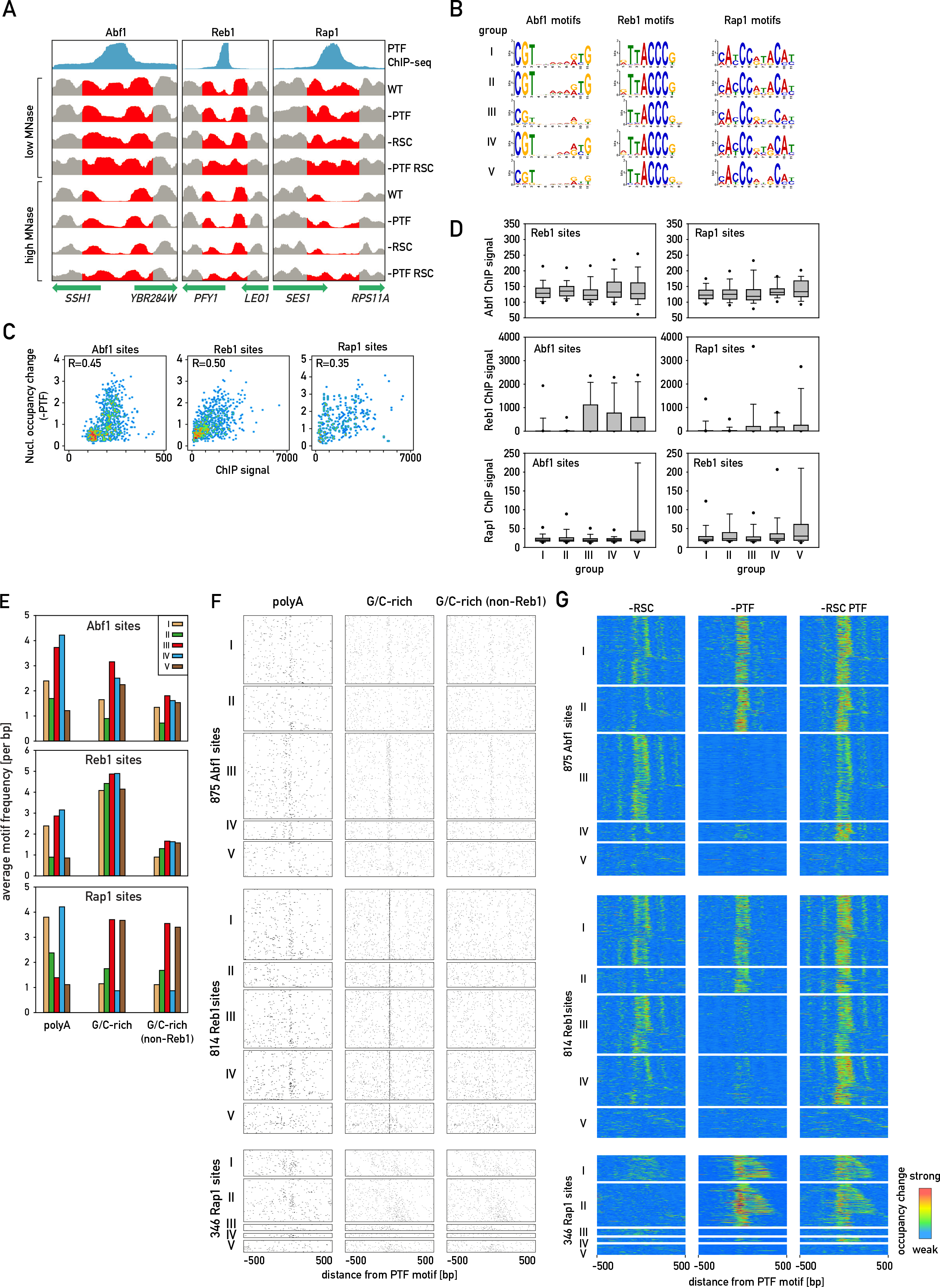
Related to Figure 5. **(A)** Nucleosome occupancy and PTF binding tracks showing examples of enhanced changes upon depletion of Sth1 and PTF comparing to depletion of just one factor alone. **(B)** Logos of consensus motifs found in each group (I-V) of PTF binding sites. **(C)** Scatterplots showing relationship between PTF binding signal in 200 bp-wide region centered on PTF binding site and nucleosome occupancy change in that region; Pearson R values are given. (D) ChIP signals of PTFs in each group of PTF binding sites. **(E)** Bar plots showing average frequency (per bp) of the following motifs: polyA, G/C-rich and G/C-rich not overlapping with Reb1 site, within 100 bp from binding sites of each PTF divided in groups shown in Figure 5B. **(F)** Heatmap showing the distribution of motifs in regions presented in Figure 5B. **(G)** Nucleosome occupancy change at PTF motif-centered regions shown as in Figure 5B but oriented and sorted with respect to closest TSS found downstream from each motif.

**Figure S6.**
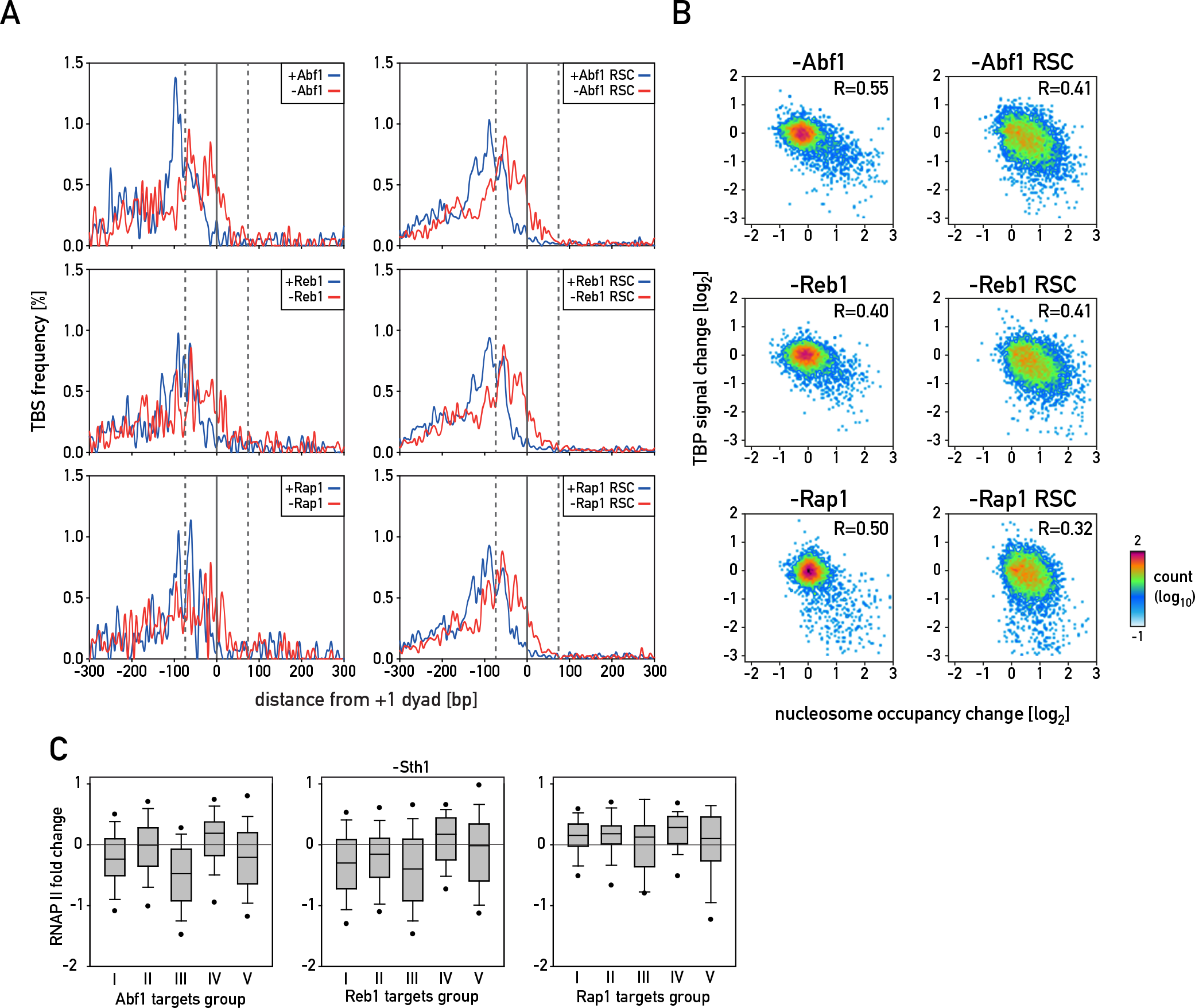
Related to Figure 6. **(A)** Plots showing frequency of TBS centered on +1 nucleosome of genes whose +1 nucleosome dyad position changed by at least 25 bp upon depletion of Abf1, Reb1 or Rap1 (plots on the left) or each PTF in combination with RSC (right), plots for wild-type +1 position in blue and position in the absence of depleted factor(s) in red. **(B)** Scatterplots showing the relationship between absolute nucleosome occupancy change and change in TBP ChIP-seq signal upon depletion of Abf1, Reb1 or Rap1 calculated in 200 bp-wide regions centered on all identified TBSs. **(C)** Boxplots showing change in RNAP II ChIP-seq signal upon depletion of Sth1 calculated for genes present in each motif group shown in Figure 5B.

